# Characterizing the power spectrum dynamics of the NREM to REM sleep transition

**DOI:** 10.1101/2023.06.14.544943

**Authors:** Diego Serantes, Matías Cavelli, Joaquín González, Alejandra Mondino, Luciana Benedetto, Pablo Torterolo

**Affiliations:** Departamento de Fisiología, Facultad de Medicina, Universidad de la República, Montevideo 11800, Uruguay; Department of Psychiatry, University of Wisconsin-Madison, Madison, Wisconsin 53719, USA; Brain Institute, Federal University of Rio Grande do Norte, Natal 59078, Brazil; Departamento de Clínicas y Hospital Veterinario, Unidad de Medicina de Pequeños Animales, Neurología, Universidad de la República, Montevideo 13000, Uruguay; Department of Clinical Sciences, College of Veterinary Medicine, North Carolina State University, North Carolina 27695, USA

**Keywords:** intermediate stage, slow wave sleep, REM sleep, EEG, sleep spindles

## Abstract

The transition from NREM to REM sleep is considered a transitional or intermediate stage (IS), characterized by high amplitude spindles in the frontal cortex and theta activity in the occipital cortex. Early reports in rats showed an IS lasting from 1 to 5 s, but recent studies suggested a longer duration of this stage. To further characterize the IS, we analyzed its spectral characteristics on electrocorticogram (ECoG) recordings of the olfactory bulb (OB), motor (M1), somato-sensory (S1) and secondary visual cortex (V2) in twelve Wistar male adult rats. By comparing the IS to consolidated NREM/REM epochs, our results reveal that the IS has specific power spectral patterns that statistically differ from both NREM and REM sleep states. Specifically, the main findings were that sigma (11-16 Hz) and beta (17-30 Hz) power in OB, M1, and S1 increased during the IS compared to NREM and REM sleep and began 55 s before REM sleep onset. Additionally, low gamma (31-48 Hz) in the OB started transitioning from NREM levels to REM ones 65 s before its onset. Finally, the high-frequency oscillations (102-198 Hz) in OB, M1, and S1 showed a power increase that began 40 s before REM sleep and reached REM sleep values 10 s after its onset. Thus, we argue that the NREM to REM transition contains its own spectral profile and is more extended than previously described.

## 1. Introduction

Sleep is a complex and dynamic process that is composed of two major states, non-rapid eye movement (NREM) sleep and rapid eye movement (REM) sleep (Siegel, 2008; Vanini & Torterolo, 2021). In mammals, polysomnography is the standard method used to discriminate these states, which simultaneously measures the electrocortical and electromyogram activity (Carskadon & Dement, 2005; Torterolo et al., 2022). In rats, NREM sleep is characterized by cortical high voltage and slow oscillations with a predominance of delta frequencies (0.5 - 4 Hz) and sleep spindles or sigma band activity (11 - 16 Hz) (Mondino et al., 2020; Mondino, Torterolo, et al., 2022; Sullivan et al., 2014), while in REM sleep the electrocorticogram (ECoG) shows low voltage and high-frequency oscillations that are nested in regular theta activity (5 - 9 Hz) (Cavelli et al., 2018; González et al., 2020; Mondino et al., 2020; Mondino, Torterolo, et al., 2022). Between these two states, classical studies described a short (1 - 5 s) transitional state, known as Intermediate Stage (IS) (Gottesmann, 1973), characterized by simultaneous high amplitude spindles in the frontal cortex and theta activity in occipital areas (Gottesmann, 1973; Gottesmann et al., 1998; User et al., 1980).

Despite early efforts suggesting the IS is a NREM-REM mixture, this stage’s short duration and scarcity limited our understanding of its characteristics. However, recent reports gained new insights into this transitional period by studying the dynamics of the high-frequency oscillations, reporting a duration of up to 20 s (Sánchez-López et al., 2018). Nevertheless, while this method precisely delimited REM onset, it is still unknown how the transition emerges during NREM sleep and if this transition constitutes a proper sleep state.

Here, by comparing the IS spectral characteristics with consolidated NREM and REM states, we were able to pinpoint its specific attributes, allowing us to unveil a much longer transition than previously reported and suggesting that the IS is not simply a mixture of NREM and REM activity.

## 2. Methods

### 2.1. Experimental animals

Twelve Wistar male rats (250-300g) were used for this study. All procedures were carried out in agreement with the National Animal Care Law (18611) and with the ‘‘Guide to the care and use of laboratory animals” (8th Edition, National Academy Press, Washington DC, 2010), and approved by the Institutional Animal Care Committee (Exp. No 070153–000332-16). Animals were maintained on a 12-h light/dark cycle (lights on at 6.00 AM) with unrestricted access to food and water. Special efforts were taken to minimize the pain, discomfort, or stress of the animals.

### 2.2. Surgical procedures

The animals were chronically implanted with intracranial cortical electrodes for polysomnography (ECoG and electromyogram, EMG) recordings. Similar procedures were employed as in previous studies (Cavelli et al., 2017; González et al., 2019, 2022; Mondino, González, et al., 2022). Anesthesia was induced with a mixture of ketamine-xylazine (90 mg/kg and 5 mg/kg i.p., respectively). Rats were positioned in a stereotaxic frame and the skull was exposed. As shown in **Figure 1A**, stainless steel screw electrodes were fixed in the skull according to Paxinos and Watson (2013) atlas (Paxinos & Watson, 2013). All electrodes were placed in the right hemisphere, and separated from each other by 5 mm. The electrodes were positioned above the olfactory bulb (OB, L: +1.25 mm, AP: +7.5 mm), motor cortex (M1, L: +2.5 mm, AP: +2.5 mm), somatosensory cortex (S1, L: +2.5 mm, AP: -2.5 mm) and secondary visual cortex (V2, L: +2.5 mm, AP: -7.5 mm). A reference electrode was placed over the cerebellum (Cer, L: 0 mm, AP: -11 mm). To record the EMG, two electrodes were inserted into the neck muscles. The electrodes were soldered into a socket and fixed to the skull with dental acrylic cement. At the end of the surgical procedures, an analgesic (ketoprofen, 1 mg/kg, s.c.) was administered.

**Figure 1.**
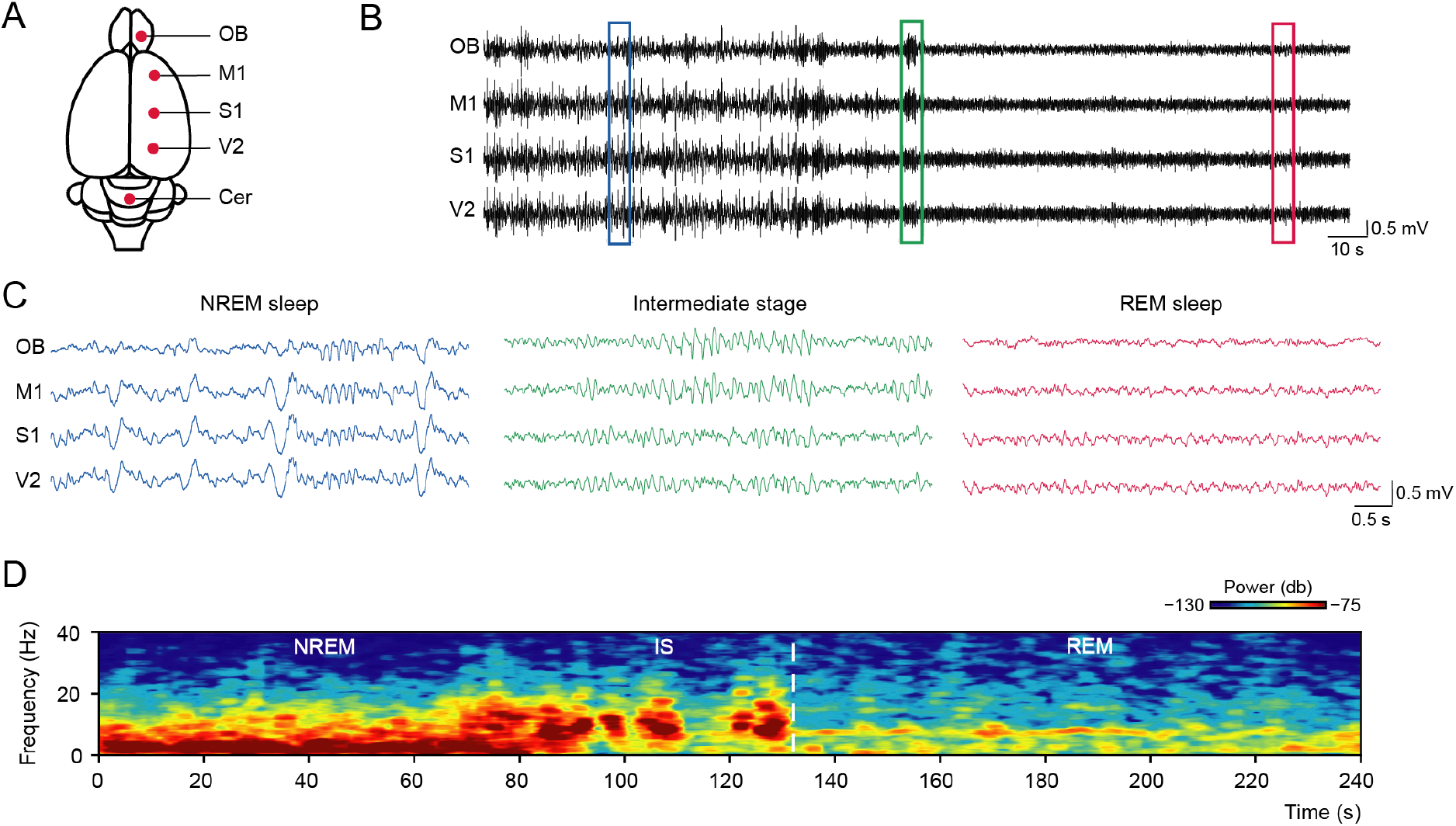
The IS shows a distinctive ECoG pattern and spectral dynamics compared to both NREM and REM sleep. **A** Schematic representation of the placement of the recording electrodes. OB: olfactory bulb; M1: primary motor cortex; S1: primary somatosensory cortex; V2: secondary visual cortex; Cer: cerebellum. **B** Raw recordings during a transition from NREM to REM sleep in a representative animal. Color rectangles indicate the zoomed segment of each sleep state. **C** Zoomed representative segments (5 s of duration) of each sleep state, showing their ECoG characteristics. **D** Representative time-frequency spectrogram of a transition from NREM to REM sleep in the M1 cortex. The vertical dashed line marks the last spindle occurrence. It is notable the distinctive spectral pattern of the IS.

### 2.3. Experimental sessions

The animals were housed in acrylic cages containing wood shaving material in a temperature-controlled room (21−24 °C), having food and water *ad libitum*. Experiments were conducted under the light period (9:00 AM to 3:00 PM) on a 12-h light/dark cycle (lights on at 6 AM). The recordings were performed using a rotating commutator to allow the rats to move freely within the recording box. Bioelectric signals were amplified (×1000), filtered (0.1–500 Hz), sampled (1024 Hz, 16 bits), and stored in a PC using Spike 2 software version 9.04 (Cambridge Electronic Design, Cambridge, UK).

### 2.4. Data analysis

We determined waking and sleep states using 10-second epochs according to standard criteria (Cavelli et al., 2015). In order to characterize the IS, we selected four effective (non-abortive) transition periods from each animal. For each transition, we selected a 160-second time-window that included the last 80 s of NREM sleep and the first 80 s of the following REM sleep episode. We delimited the end of the NREM sleep period (and the beginning of REM sleep) with the last spindle occurrence. In addition, we selected four 80-second segments of consolidated NREM and REM sleep for each animal, which we later used to compare transition characteristics. In summary, the whole data base consisted of 48 NREM to REM transitions (160-second windows), and 48 consolidated NREM and REM sleep episodes (80-second windows).

Similar to our previous works (Cavelli et al., 2017; González et al., 2021), we employed multitaper spectrograms to transform each ECoG segment into its time-frequency representation (Prerau et al., 2017) (available for python at https://prerau.bwh.harvard.edu). We created one spectrogram for each sleep period (i.e., four 160-second periods of transition and four 80-second periods of NREM and REM sleep) and then averaged them to obtain three mean spectrograms per animal (i.e., one for NREM sleep, one for the transition period, and another for REM sleep). The ECoG spectrograms were generated with a sampling frequency of 1024 Hz, a frequency range of 0.5 - 512 Hz, a window size of 4 s, 6 tapers, and a time-bandwidth product of 4. All line noise artifacts (50 Hz) were eliminated. To align the transition spectrograms, we used the end of the last spindle as a reference point. Next, we developed a Python script to compare each time-frequency bin of the transition spectrogram to the spectral power distribution for that specific frequency along the 80-second NREM and REM sleep periods. If a time-frequency bin fell outside the confidence interval (confidence level of 95%) of both NREM and REM density distributions, we considered it significant and assigned a value of 1 to that time-frequency bin; otherwise, we assigned a value of 0. The workflow employed for this analysis is shown in **Figure 2**.

**Figure 2.**
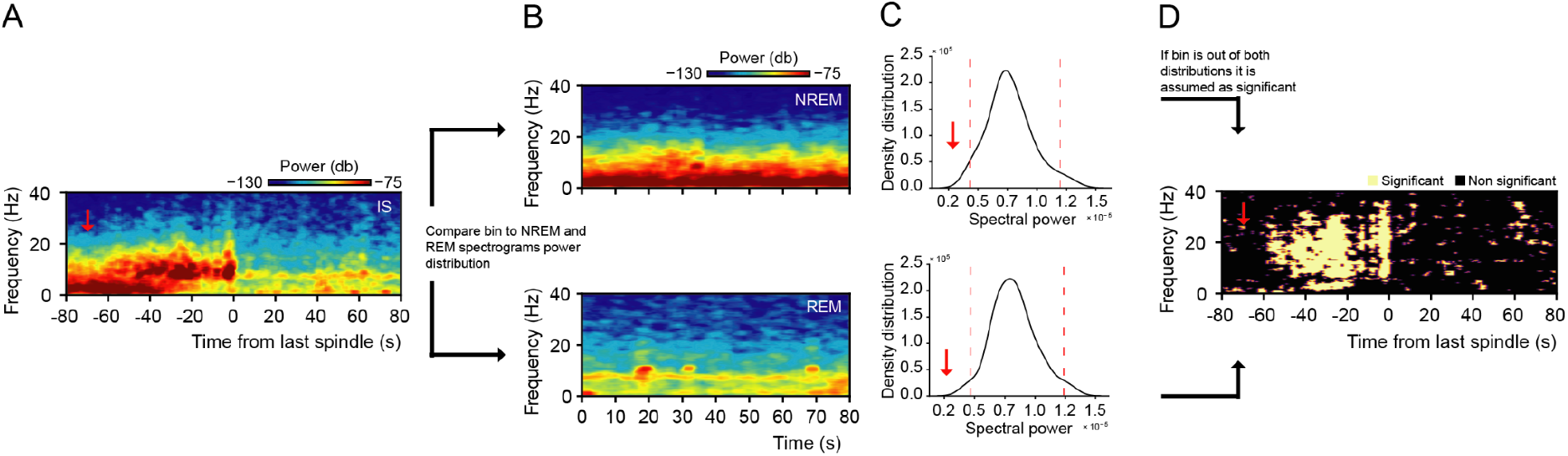
Method used for discrimination of significant time-frequency bins. **A** Averaged spectrogram from the M1 cortex during the transition from NREM to REM sleep in a representative animal (average of 4 spectrograms). The x-axis zero corresponds to the last spindle used for spectrogram alignment prior to averaging. The red arrow indicates the time-frequency bin analyzed in this example. **B** Averaged spectrograms of NREM sleep (top) and REM sleep (bottom) employed for comparison with the analyzed time-frequency bin (average of 4 spectrograms per state from the same animal). **C** Spectral power density distribution along the 80 s of the frequency being analyzed (in this example, we are analyzing the 24.5 Hz frequency). Red dotted lines indicate the 2.5th and 97.5th percentiles, respectively. The red arrow signifies that the analyzed time-frequency bin (same as marked in **A**) lies outside the 95% confidence interval of the NREM and REM sleep density distributions for the given frequency. Thus, it is considered a significant time-frequency bin. These distributions were constructed with the data of all the NREM or REM sleep spectrograms of the same animal. **D** “Difference spectrogram” showing which values from the spectrogram in **A** exhibit significant differences compared to both NREM and REM sleep. White-colored time-frequency bins represent significant differences, while black-colored ones do not. Red arrow marks the time-frequency bin being analyzed. Note that since the bin was outside both distributions confidence intervals, the dot is marked as white.

For the calculation of the absolute and relative power spectrum (i.e., relative to the total power of the signal) of the ECoG, we employed the ‘Welch’ method (welch function, available at the scipy library, Python 3.9, parameters: sampling frequency = 1024; nperseg = 1024; Window = Hann window). Frequency bands were defined as: delta (0.5 - 4 Hz); theta (5 - 9 Hz); sigma (11 - 16 Hz); beta (17-30 Hz); low gamma (31-48 Hz); high gamma (52-98 Hz); and high-frequency oscillations (HFO, 102-198 Hz). We subdivided the transition period into 5-second segments, and we calculated the power spectrum for each segment. The same procedure was made for each animal and averaged together. For NREM and REM sleep, we calculated the power spectrum of the time windows used before (four windows per animal) and then averaged them. Repeated measures ANOVA was performed to compare the band power of each 5-second segment of the transition to the band power of the stationary segments of NREM and REM sleep. In cases where the ANOVA indicated significant differences, *post hoc* comparisons were performed using a t-test between the transition and both NREM and REM sleep. Statistical significance was set at p ≤ 0.01. Only the transition values that demonstrated *post hoc* significance with both NREM and REM sleep were considered significant. To ensure a more stringent time delimitation of the transition, we only considered two consecutive significant values as a time starting point.

## 3. Results

### 3.1. The IS has a unique power spectrum profile

**Figure 1B-C** shows a representative recording of the NREM to REM transition. Notice that during the IS, the sleep spindles are intermixed with theta activity, consistent with previous findings (Gottesmann, 1973). Then, we analyzed the power spectrum dynamics of the IS and compared it to those of NREM and REM sleep. To this end, we generated time-frequency spectrograms; an example of a NREM to REM transition is shown in **Figure 1D**. This analysis shows that the power spectrum of the IS changes 40 to 50 s before the beginning of REM sleep.

Next, we investigated whether the observed changes in the power spectrum resulted from the attenuation of NREM and the onset of REM sleep (i.e., a hybrid state) or if there were spectral manifestations that were unique to the IS. To assess this, we analyzed the spectrograms of the IS in 12 rats using the method described in **Figure 2**. Briefly, we located transition periods by tracking the last sleep spindle before REM onset. Next, we computed spectrograms surrounding this trigger (**Fig. 2A**) and during stable NREM and REM episodes (**Fig. 2B**). Then, we constructed the distributions over time of each frequency component for the stable NREM and REM spectrograms (**Fig. 2C**). Finally, we compare each time-frequency bin of the transition spectrogram with its corresponding NREM-REM frequency distribution. All spectrogram points below the 2.5th or above the 97.5th percentile for both NREM and REM distributions were considered significant and assigned a 1 (nonsignificant points a 0). Thus, by developing this method, we obtained a new time-frequency spectrogram of the transition period, which indicates which frequency components are unique to this stage (**Fig. 2D**). It should be noted that our construction is able to tell us which time-frequency bins statistically differ from both NREM-REM distributions, while the sign of the change (i.e., increase/decrease) can be then inferred from the normal spectrogram.

Figure 3 shows the comparison between the average transition spectrogram (Panel A, averaged across animals) and our modified “difference” spectrogram (Panel B). If we look at the average transition spectrogram, we can observe an increase in the sigma band power, simultaneously accompanied by a decrease in the delta band power, starting approximately 20 s before the onset of REM sleep. These changes varied across the different brain regions, suggesting that the power spectrum dynamics of the IS are heterogeneous. In contrast, **Figure 3B** shows a much broader transition period (the color gradient from purple to white represents the time-frequency bins that were significant in a percentage of the animals). Specifically, we can observe that the increase in sigma activity (even spanning to the beta band) starts uniformly in all regions approximately 50 s prior to REM sleep onset. Interestingly, our method shows that the delta band power decrease occurred after the sigma increase, approximately 30 s before REM onset. Since no consistent significant changes were observed above 40 Hz, these frequencies were not plotted. Thus, these results show that the IS has its own spectral signature and dynamics, involving a precise sigma-delta coordination.

**Figure 3.**
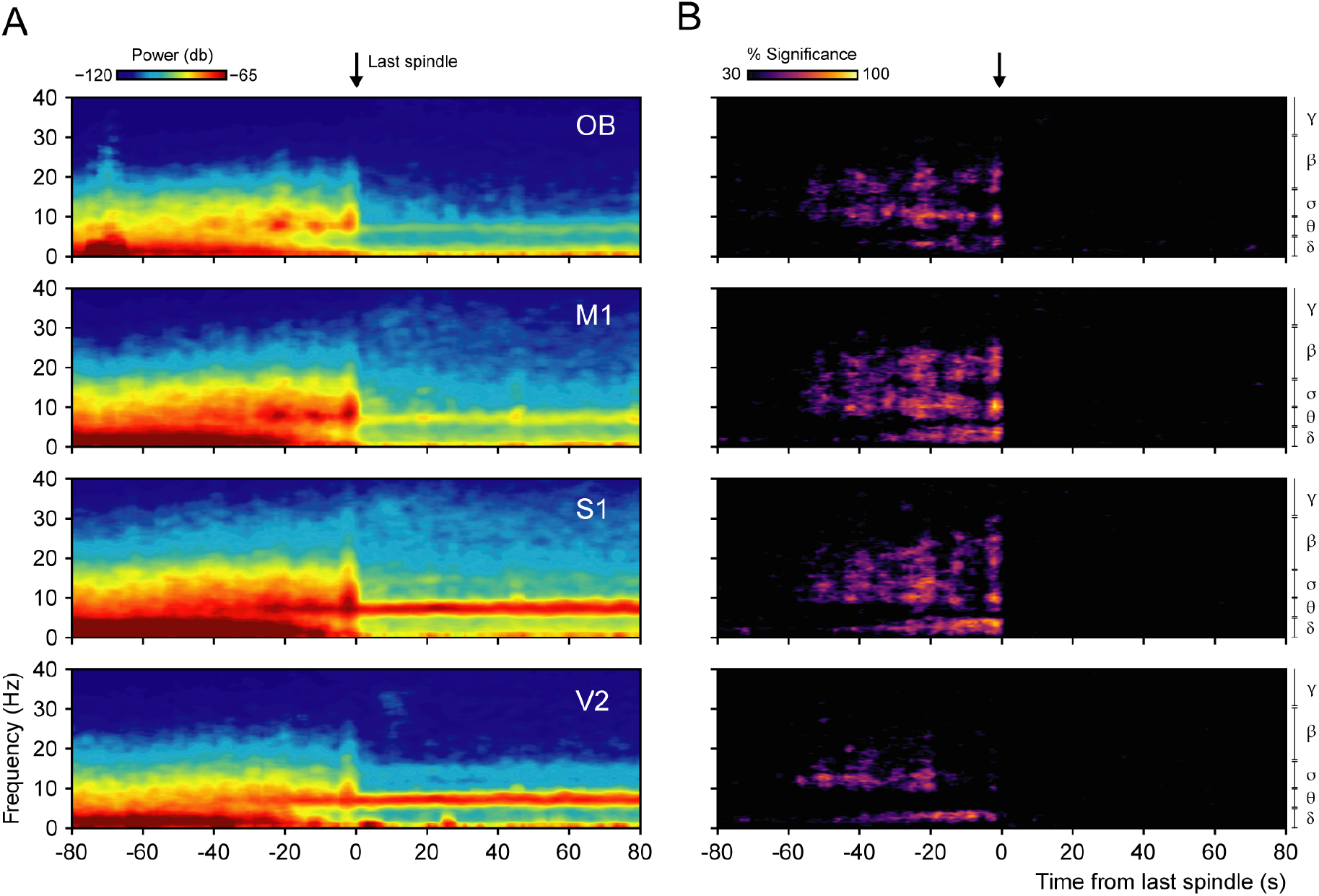
The intermediate stage shows significant power spectrum differences from both NREM and REM sleep. **A** Averaged spectrograms from all animals (n=12) showing absolute power dynamics in the frequency range of 0 - 40 Hz. The zero point indicates the end of the last spindle prior to REM sleep onset, serving as the alignment reference for averaging the spectrograms. **B** Averaged “difference” spectrograms showing which values of the spectrograms in **A** are significantly different from both NREM and REM sleep. The statistical method employed is outlined in **Figure 2**. The frequency bands analyzed are indicated on the right side, including delta *δ* (0.5 - 4 Hz); theta *θ* (5 - 9 Hz); sigma *σ* (11 - 16 Hz); beta *β* (17 - 30 Hz); low gamma *γ* (31 - 40 Hz). Black arrows in **A** and **B** mark the end of the last spindle preceding the onset of REM sleep.

### 3.2. Temporal dynamics of the lower ECoG frequency bands in the IS

To confirm our results, we employed a complementary approach to measure the temporal dynamics of the IS. We binned the transition in separate 5 s windows, calculated the power spectrum, and averaged the different frequency bands, thus reducing the variability inherent to the high-frequency resolution spectrograms. **Figure 4** shows the averaged power changes of the IS for the lower frequency bands (delta, theta, and sigma) in intervals of 5 s. These traces were then statistically compared to the average power of NREM and REM sleep stationary segments.

**Figure 4.**
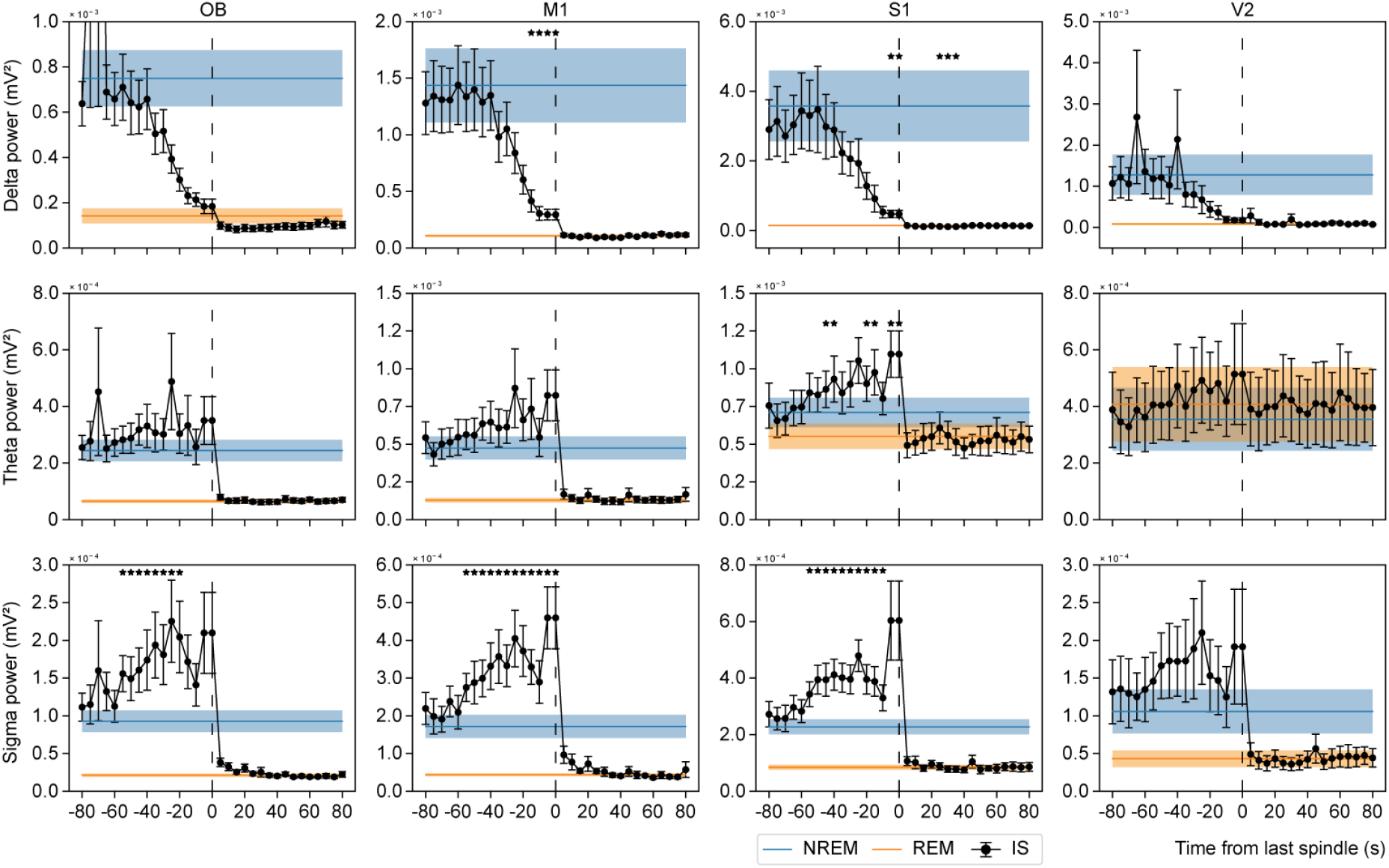
The IS has region- and frequency-specific onset and offset times. The plots show the average power dynamics of the IS for delta (0.5 - 4 Hz), theta (5 - 9 Hz), and sigma (11 - 16 Hz) bands, categorized by cerebral area. Power spectrum was calculated using Welch’s method, with 5-second segments used for each data point. The points represent the mean power of the population (n=12), and the T-bars indicate the standard error of the mean. The line representing NREM and REM sleep displays the mean power of the population for an 80-second duration segment, while the shaded line depicts the standard error of the mean. Notice that the NREM and REM sleep power spectrum appears as a line because it was calculated over the entire 80-second segment. In contrast, for the transition, the Welch method was applied to 5-second segments. Significant data points highlight the elements that differ significantly from both NREM and REM sleep. Only consecutive (two or more) points are illustrated.

Consistent with our spectrogram results, we observed an increase in sigma band activity within the OB, M1, and S1 relative to stationary NREM and REM values, 55 s prior to the occurrence of the last spindle (OB: F(2, 22) = 40.5, p < 0.001; M1: F(2, 22) = 41.8, p < 0.001; S1: F(2, 22) = 43.0, p < 0.001). Subsequently, a sudden drop in power to REM values was observed. Conversely, no significant changes were detected in V2 despite exhibiting a similar profile. Additionally, in M1, a decrease in delta band power was observed 15 s prior to REM sleep (F(2, 22) = 14.7, p < 0.001), while in S1, the decrease occurred at 5 s (F(2, 22) = 11.4, p < 0.001). Note that we also observed a small significant difference in S1 delta after REM onset; however, this is due to a slight decrease compared to the low error levels of REM delta. Moreover, no significant changes in delta band power were noted in OB and V2. In regard to the theta band, no significant changes were observed, except in S1, where intermittent significant changes initiated 45 s before REM sleep (F(2, 22) = 8.0, p = 0.002). The limited significant findings for delta and theta bands could potentially be attributed to the high variability observed among subjects, a factor that was attenuated when analyzing the relative power (i.e., relative to the total power) as depicted in **Supplementary Figure S1**.

### 3.3. Temporal dynamics of the higher ECoG frequency bands in the IS

Finally, we complemented our results by studying the temporal dynamics of the higher frequency bands, employing the same method shown in **Figure 4**. The average power of the IS within the higher frequency bands (beta, low gamma, high gamma, and HFO) is presented in **Figure 5**. Interestingly, the beta band exhibited a pattern of change similar to that of the sigma band (**Figure 3**). Additionally, significant changes in the beta band started 55 s before REM sleep onset in the OB and M1 regions (OB: F(2, 22) = 18.9, p < 0.001; M1: F(2, 22) = 34.2, p < 0.001), and 45 s in S1 (F(2, 22) = 15.2, p < 0.001), providing further support for the last observation since the beta band changes start at the same time as sigma band.

**Figure 5.**
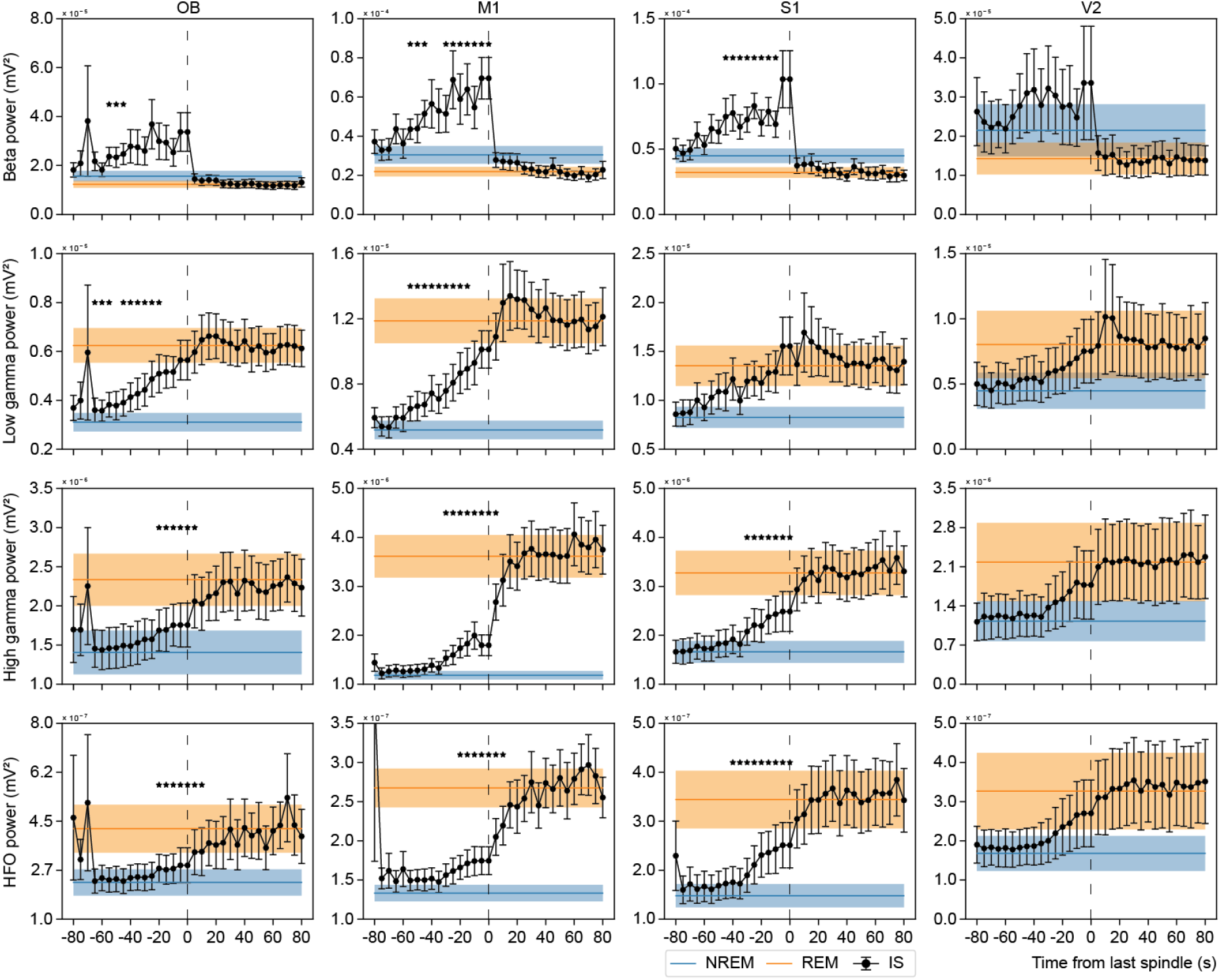
The IS region- and frequency-specific onset and offset times are also observed in the high-frequency bands. The plots show the average power dynamics of the IS for beta (17 - 30 Hz), low gamma (31 - 48 Hz), high gamma (52 - 98 Hz), and HFO (102 - 198 Hz). Power spectrum was calculated using Welch’s method, with 5-second segments used for each data point. The points represent the mean power of the population (n=12), and the T-bars indicate the standard error of the mean. The line representing NREM and REM sleep displays the mean power of the population for an 80-second duration segment, while the shaded line depicts the standard error of the mean. Significant data points highlight the elements that differ significantly from both NREM and REM sleep. Only consecutive (two or more) points are illustrated.

On the other hand, excluding the beta band, all higher frequency bands (low-gamma, high-gamma, and HFO) show a common cortex-wide profile consisting of an increase to a REM plateau value. Specifically, significant changes in the low gamma band began 65 s before REM sleep in the OB (F(2, 22) = 56.9, p < 0.001), and 55 s in the M1 cortex (F(2, 22) = 42.2, p < 0.001), while no significant changes were observed in S1 and V2.

Regarding the power of the high gamma and HFO bands, the significant changes extended beyond the last spindle occurrence (**Figure 5**). Notably, in the high gamma band, changes initiated 30 s before REM sleep onset in M1 and S1 (M1: F(2, 22) = 41.2, p < 0.001; S1: F(2, 22) = 35.0, p < 0.001), and 20 s in the OB (F(2, 22) = 34.7, p < 0.001). Moreover, in OB and M1, the significant changes persisted for 5 s after the last spindle. Furthermore, the HFO band exhibited significant changes in S1 beginning 40 s before REM sleep onset (F(2, 22) = 27.8, p < 0.001), and in OB and M1, 20 s before REM (OB: F(2, 22) = 30.5, p < 0.001; M1: F(2, 22) = 43.8, p < 0.001), which continued for 10 s after the last spindle occurrence. No significant values were observed in V2 for any of the higher frequency bands, likely due to the substantial variability between animals.

## 4. Discussion

In the present study, we analyzed the whole power spectrum of the ECoG of the rat along the cortex’s anterior-posterior axis with high temporal resolution. Our results suggest that the IS in the rat significantly differs from the ECoG hallmarks which characterize NREM and REM sleep (Carskadon & Dement, 2005; Gervasoni et al., 2004; Mondino et al., 2020; Mondino, Torterolo, et al., 2022; Nir et al., 2011; Steriade, 2000). These significant spectral changes start 65 s before the onset of REM sleep and continue for 10 s after the last spindle occurrence, as summarized in **Figure 6**; note the region- and frequency-specific onset and offset times for the IS among different brain regions and frequency bands. Notably, our results show a much longer intermediate stage than previously described (Gottesmann, 1973; Gottesmann et al., 1998; Sánchez-López et al., 2018). Thus, we argue that the IS is not a hybrid state (i.e., simply a transition from NREM to REM sleep) but rather a unique sleep state with its own power spectrum profile and, thus, its unique network dynamics.

**Figure 6.**
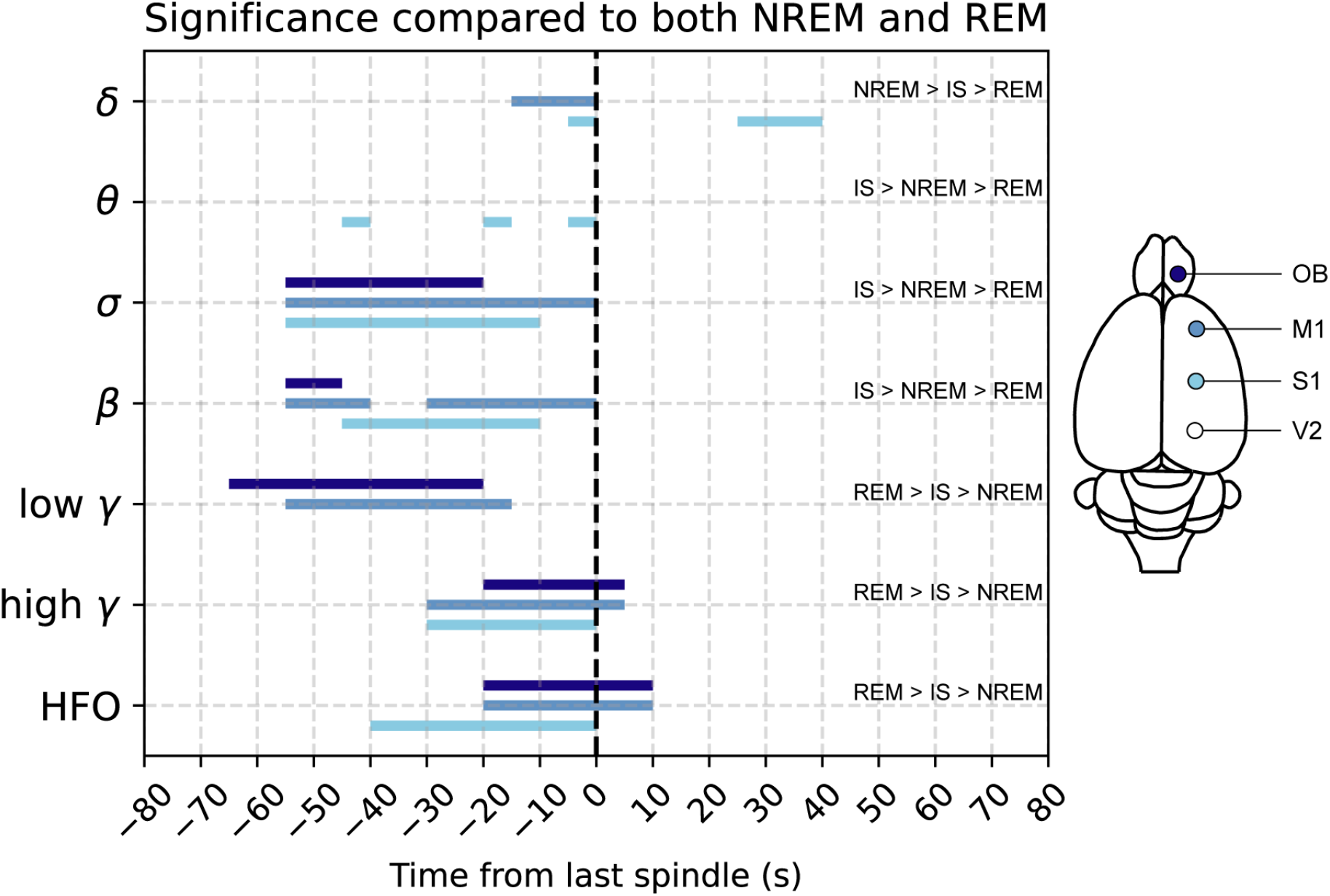
The IS begins 65 s before REM sleep onset. Bars show the time in which the power of each band is significantly different (p ≤ 0.01) compared to both NREM and REM sleep for all 12 animals analyzed. The color code (right) indicates each area represented (notice that V2 is not represented because of its non-significant changes). The dashed line indicates the last spindle occurrence before REM sleep onset.

### 4.1 Temporal dynamics of the IS

Our study shows a significant increase in sigma band power (that corresponds to the spindle frequency) prior to the onset of REM sleep, consistent with previous findings (Bandarabadi et al., 2020; De Gennaro & Ferrara, 2003). In fact, enhancing the activity of thalamic reticular nucleus cells or their projection to other thalamic nuclei increase the spindle rate and transitions to REM sleep (Bandarabadi et al., 2020). Moreover, as REM sleep commences, the spindles abruptly disappear, confirming early observations (Gottesmann, 1992). Additionally, sigma band power increases well before the decrease in delta power, revealing a novel aspect of the well-established inverse relationship between sigma and delta bands (Bonjean et al., 2011; De Gennaro et al., 2000; Werth et al., 1997). This interplay becomes even more apparent when examining the relative power spectrum of the delta and sigma bands (**Supplementary Figure S1**). Recent evidence shows that active inhibition generated by somatostatin-positive (SOM+) cells is directly related to the slow wave’s onset (Funk et al., 2017). Furthermore, both the decrease in the firing of these cells and their chemogenetic inhibition decrease delta and increase sigma bands (Funk et al., 2017; Niethard et al., 2018; Spano et al., 2022). In that sense, our results suggest that the IS could represent a disinhibited state with a progressive withdrawal of SOM+ tone. Finally, in concordance with our temporal dynamics results, studies conducted in the Locus Coeruleus showed a decrease in noradrenaline levels and spiking activity that began ∼ 40 s before REM sleep onset (Aston-Jones & Bloom, 1981; Kjaerby et al., 2022; Osorio-Forero et al., 2021, 2022).

The beta band power exhibits a remarkably similar pattern of change to the sigma band (**Figure 5**). We attribute this similarity to spindles increasing their frequency up to the beta range. These results highlight the need for circuit-level definitions when delimiting frequency bands (Fernandez-Ruiz et al., 2023; Lopes-Dos-Santos et al., 2018), indicating that the same neural circuit oscillates between 11 and 30 Hz during the IS. Moreover, in studies conducted on rats, the NREM sigma power displays a shoulder (Fernandez & Lüthi, 2020; Mölle et al., 2009), suggesting a broad frequency range. Similarly, in mice, the NREM sleep power spectrum exhibits an increase in the 9 - 25 Hz range prior to REM sleep onset (Astori et al., 2011; Cueni et al., 2008). Collectively, these pieces of evidence lead us to interpret the sigma and beta bands of the IS as a unified entity whose augment is the most relevant signature of this sleep state.

Regarding the gamma band (low and high gamma), our results indicate that low gamma power increases concurrently with sigma band power 55 s prior to REM sleep onset, while the increase in high gamma band power is delayed but lasts for an additional 5 s compared to the differences observed in the sigma band (**Figure 6**). Consistent with previous reports (Ayoub et al., 2012; Weber et al., 2021), we observed an association between sleep spindles and gamma cortical activity during the IS, which relates to the mechanisms of memory consolidation (Ayoub et al., 2012; Weber et al., 2021). Finally, in agreement with our results, the HFO increase was previously described and employed to delineate the NREM to REM transition (Sánchez-López et al., 2018).

### 4.2 IS in other species

The IS has been previously described in humans (Grubar, 1983), cats (Gottesmann et al., 1984), and mice (Glin et al., 1991). Specifically in cats, REM sleep is always preceded by ponto-geniculo-occipital (PGO) waves (Datta, 1997; Duysan-Peyrethon et al., 1967; Jouvet et al., 1959; McCarley et al., 1978) leading to consider the PGO waves occurrence as the transitional period in this species. Remarkably, these waves also appear approximately 1 minute before the beginning of REM sleep (Ursin & Sterman, 1981), supporting the notion of an extended IS. In fact, Steriade et al. (1990) demonstrated that neurons in the mesopontine cholinergic region increase their firing rate about 1 minute before the REM sleep onset (Steriade et al., 1990). Carrera-Cañas et al. (2019) recently analyzed the IS in cats and concluded that it constitutes a state with characteristics that are distinct from both the preceding NREM sleep and the following REM sleep. The IS presented large spindles, low delta activity, and PGO waves that were similar to a state of dissociated sleep, called SPGO state, produced following the administration of carbachol into the perilocus coeruleus, a nucleus in the pontine tegmentum. Note that the caudolateral peribrachial area, considered the PGO-generator area in the cat (Datta et al., 1992), has been proposed to play a primary role in coordinating the activities of various central and peripheral systems during the transition from NREM to REM sleep (Torterolo et al., 2011).

With respect to mice, the IS described by Glin et al. (1991) had similar characteristics to those in rats, with high amplitude sleep spindles interspersed with delta waves in the frontal cortex, while theta activity was registered in the hippocampus. In addition, the IS duration was about 16 s, and the theta activity analyzed was significantly different from that of REM sleep.

Although an increase in the sigma band power previous to REM sleep onset is a classical finding (Ayoub et al., 2012; Purcell et al., 2017), there are almost not human studies with the emphasis in the IS. Grubar (1983) focused on the analysis of the IS in patients with mental deficiency, where IS occurred more frequently than in control subjects. In addition, the mentally deficient patients expressed the first IS episode of the sleep cycle with no subsequent REM sleep, while control subjects expressed all IS episodes with a subsequent REM sleep episode.

The information collected from all studies about the IS suggests that it is a general sleep phenomenon expressed in several species. The characteristics of the IS appeared to be similar between species, particularly between rats, mice, and cats, while further investigation is needed in humans to give an accurate comparison.

The functional properties of this sleep state are still unknown. However, it has been described that the thalamocortical transmission level during the IS was the lowest among all sleep stages (Gandolfo et al., 1980), resembling a “cerveau isole” preparation (Gottesmann et al., 1980). In fact, Gottesman (1996) suggested that during the IS the forebrain is disconnected from the brainstem. On the other hand, the presence of pontine (P) waves in rats (analogous to the PGO waves in other species) during IS, and the fact that learning increases the number of IS episodes (Datta, 2000, 2006), indicate that the IS could be a state with important mnemic properties. In this regard, Ramirez-Villegas et al. (2020) and Tsunematsu et al. (2020) described in monkeys and mice, respectively, the coupling of the PGO waves with hippocampal ripples and theta oscillations and proposed that this synchronization has a critical role in learning and memory (Ramirez-Villegas et al., 2021; Tsunematsu et al., 2020).

## 5. Conclusion

Our results suggest that the IS could be interpreted as an independent sleep state; i.e., a state different from NREM and REM sleep. This state in rats lasts approximately 65 s, which is longer than previously described. More studies are needed to identify the mechanisms responsible for generating this particular transition period between NREM and REM sleep and its functional relevance.

## Acknowledgments

This research was supported by the CSIC-I+D grupos 2022-group ID-22620220100148 grant, and the “Programa de Desarrollo de Ciencias Básicas”, from Uruguay.

