## Supplementary material for "Characterizing the power spectrum dynamics of the NREM to REM sleep transition"

**Figure S1**

**Figure S2**

**Figure S3**

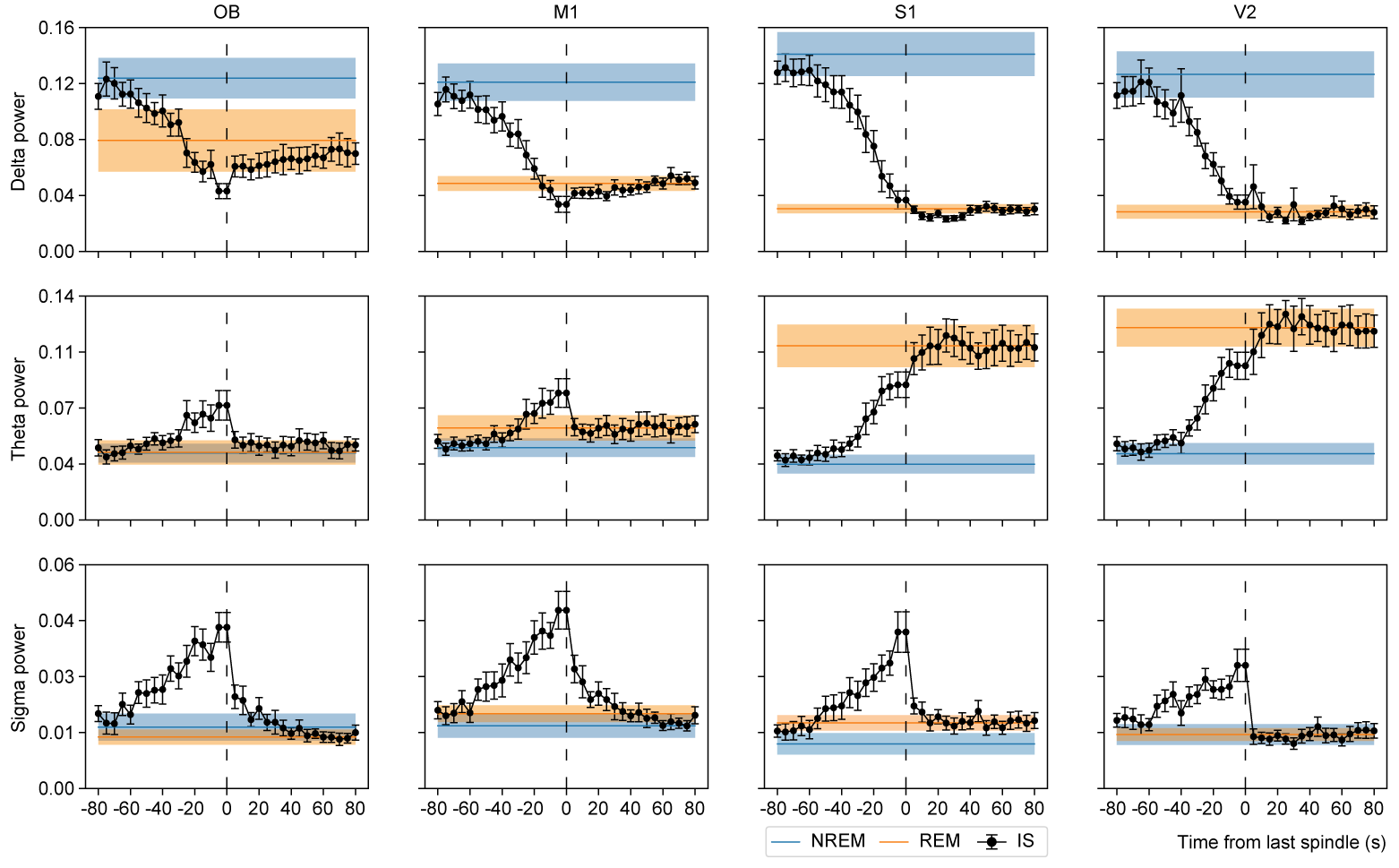

**Figure S1. Relative power dynamics of the transition from NREM to REM sleep in the lower frequency bands.**

The Figure shows the averaged relative power dynamics of the IS for delta (0.5-4 Hz), theta (5-9 Hz), and sigma (11-16 Hz) frequency bands. The relative power was calculated using the Welch method and then divided by the total power of the signal (i.e., 0 - 200 Hz). Points show the mean relative power of the population (n=12) and T bars show 2 times the standard error of the mean. The line of NREM and REM sleep shows the mean relative power of the population for a segment of 80 s of duration, and the shaded line shows 2 times the standard error of the mean. Notice that the NREM and REM power spectrum is a line because it was calculated for all the 80 s of the segment (i.e., we run the Welch method over the 80 s at once, while for the transition we run the Welch method over segments of 5 s of transition).

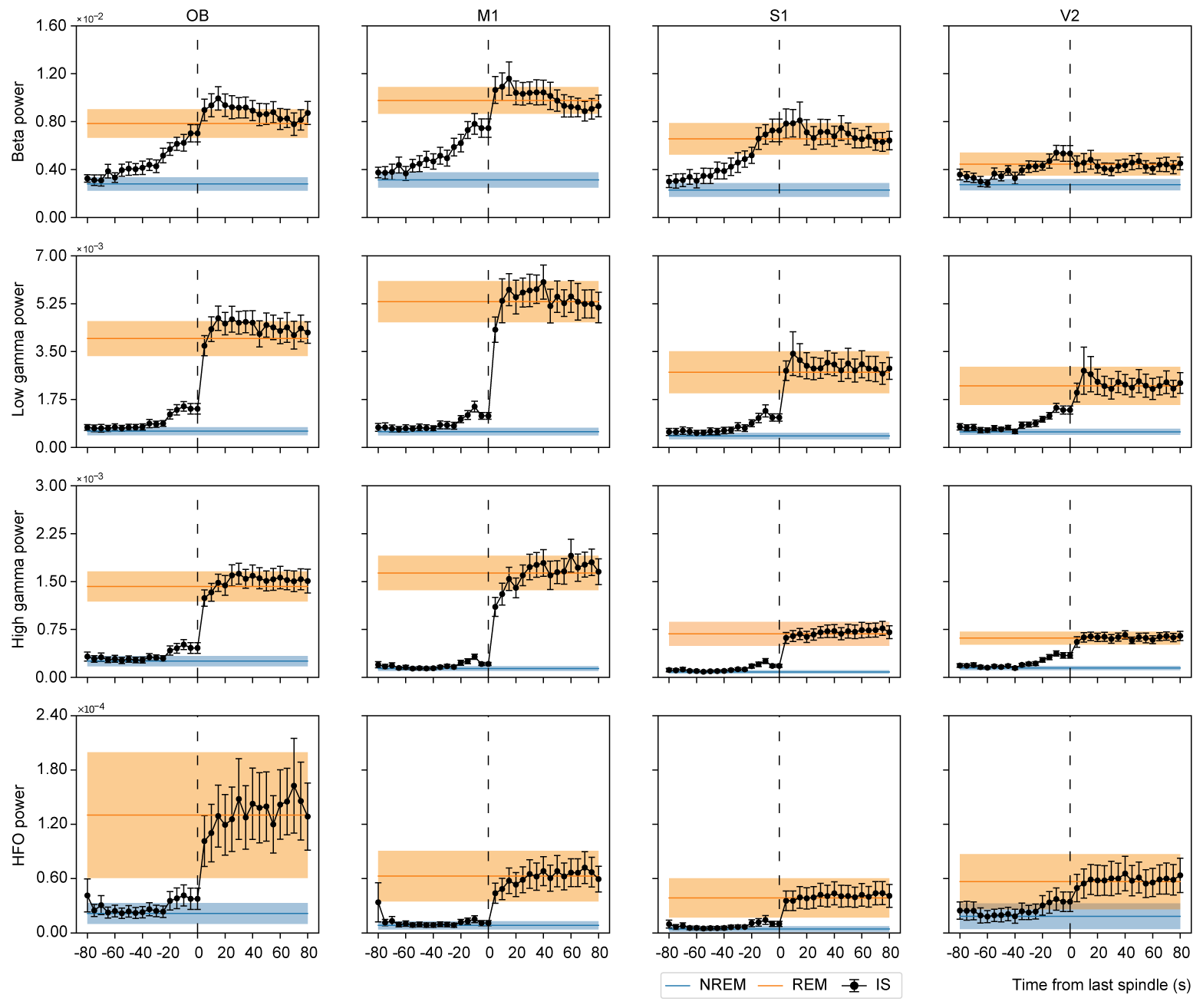

**Figure S2**. **Relative power dynamics of the transition from NREM to REM sleep in the higher frequency bands.**

Figure represents the averaged relative power of Beta (16-30 Hz), low gamma (31-48 Hz), high gamma (52-98 Hz), and HFO (102-198 Hz) frequency bands. The relative power was calculated using the Welch method and then divided by the total power of the signal (i.e., 0 - 200 Hz). Points show the mean relative power of the population (n=12) and T bars show 2 times the standard error of the mean. The line of NREM and REM sleep shows the mean relative power of the population for a segment of 80 s of duration, and the shaded line shows 2 times the standard error of the mean. Notice that the NREM and REM power spectrum is a line because it was calculated for all the 80 s of the segment (i.e., we run the Welch method over the 80 s at once, while for the transition we run the Welch method over segments of 5 s of transition).

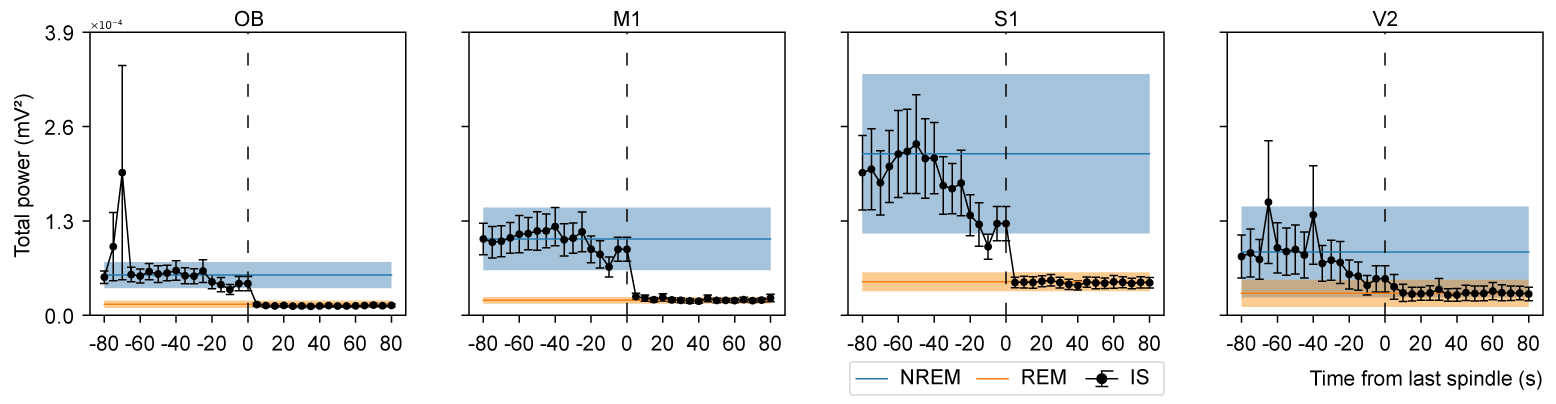

**Figure S3. Total power dynamics of the transition from NREM to REM sleep.**

The Figure represents the total power spectrum of the IS signal in the range from 0.5 - 198 Hz. The absolute power was calculated using the Welch method, tacking segments of 5 s for each dot. Points show the mean of the population (n=12) and T bars show 2 times the standard error of the mean. The line of NREM and REM shows the total power mean of the population, and the shaded line shows 2 times the standard error of the mean.
